# Harmonization of Margin of Stability Calculations and Investigation of the Impact of Foot Length, Foot Width, Gait Speed, and Body Mass

**DOI:** 10.1101/2024.09.19.613112

**Authors:** Cloé Dussault-Picard, Claire Robidou, Romain Tisserand, Yosra Cherni

**Author notes:** **Corresponding author:** Cloé Dussault-Picard.

## Abstract

The margin of stability (MoS), the minimum distance between the extrapolated center of mass and the edges of the base of support (BoS), is one of the most widely used metric to describe the mechanical stability during gait. In the current literature, the markers used to define the edges of the BoS are variable and the MoS model neglects the influence of anthropometric factors, such as foot length, foot width, and body mass. This study aimed to evaluate differences between anteroposterior (AP) and mediolateral (ML) MoS measures using various BoS edge definitions (AP: n = 3 methods, ML: n = 4 methods) and to investigate the impact of foot length, foot width, gait speed, and body mass on the MoS measures. Results show that the BoS edges definition affects the resulting MoS across the entire stance phase (AP: p<0.001 between the 3 methods; ML: p<0.001 between the 4 methods). Moreover, the AP MoS is influenced by foot length (p<0.029), as well as gait speed and body mass on both the AP (gait speed: p<0.001; body mass: p<0.038) and ML (gait speed: p<0.032; body mass: p<0.001) MoS. This study proposes a new approach based on optimal foot markers for defining the edges of the BoS, which may contribute to better assess mechanical stability during gait. Finally, the results suggest that normalizing the MoS (i.e., the AP MoS by foot length, gait speed, and body mass, and the ML MoS by gait speed and body mass) can facilitate comparisons between populations.

## 1. Introduction

Various methods have been employed to assess stability during human locomotion (Bruijn et al., 2013). These methods span from ordinal scale clinical assessments (e.g., the Berg Balance Scale (Miranda-Cantellops and Tiu, 2024) and the Functional Gait Assessment (Leddy et al., 2011)) to biomechanical measures derived from simple mechanical system (e.g., the margin of stability (MoS) (Devetak et al., 2019; Hof et al., 2005)). The MoS is one of the most widely used metric to describe the instant mechanical stability of the body configuration during pathological (Watson et al., 2021) and non-pathological gait (Ohtsu et al., 2019), and is related to the minimal impulse needed for destabilizing the walking person (Curtze et al., 2024). The MoS represents the minimum distance between the extrapolated center of mass (xCoM) and the edges of the base of support (BoS) (*Eq. 1*), and can be calculated in the antero-posterior (AP) and the medio-lateral (ML) directions of the stance phase of walking.

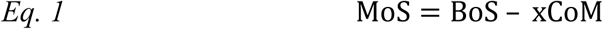

The xCOM was introduced by Hof and colleagues in 2005, and is based on the traditional linearized inverted pendulum model (Hof et al., 2005). This latter considers that the CoM location is continually changing according to even small body position changes by combining the CoM position and its velocity (*CȮM*) divided by the pendulum’s eigen frequency, i.e., the square root of gravity (9.81 m/s^2^) divided by the pendulum length (*l*) (*Eq. 2*).

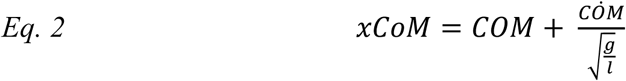

The edges of the BoS can be described using the limits of the center of pressure (CoP) (Brandt et al., 2019; Day et al., 2007; De Jong et al., 2020; Hof et al., 2007; Major et al., 2013; Van Meulen et al., 2016; Vistamehr et al., 2016) or foot markers (Beltran et al., 2014; Gates et al., 2013; Hak et al., 2015, 2014, 2013b, 2013c; Kao et al., 2014; Major et al., 2019; Martelli et al., 2017; Peebles et al., 2017, 2016; Rijken et al., 2015; Simon et al., 2017; Tisserand et al., 2018). Using foot markers, previous studies have described the lateral edge of the BoS with the fifth metatarsal marker (Beltran et al., 2014; Gates et al., 2013; Kao et al., 2014; Major et al., 2019; Martelli et al., 2017; Peebles et al., 2017, 2016), the lateral malleolar marker (Hak et al., 2015, 2014, 2013b, 2013c; Rijken et al., 2015), or the mid-point between the fifth metatarsal and the lateral malleolar markers (Simon et al., 2017; Tisserand et al., 2018). Although the definition of the AP edges seem to be more standardized, based on the toe (anterior edge) (Carty et al., 2011; Hak et al., 2013c; Kao et al., 2014; Ma et al., 2021; McAndrew Young et al., 2012; McAndrew Young and Dingwell, 2012; McCrum et al., 2014; Peebles et al., 2017, 2016; Sangeux et al., 2024; Sivakumaran et al., 2018; Tracy et al., 2019) or heel landmark (posterior edge) (Arora et al., 2019; Hak et al., 2015, 2013a; Herssens et al., 2020; Rijken et al., 2015; Wang et al., 2024; Yamaguchi and Masani, 2022), the heterogeneous methodologies relating the definition of the lateral edge of the BoS is a key factor making interpretation and comparison of MoS results challenging (Watson et al., 2021). Uncertainty persists regarding the optimal marker to define the edges of the BoS at different instants of the stance phase (Curtze et al., 2024). For the ML BoS, it can be suggested that using the midpoint on the virtual line relating the fifth metatarsal and the lateral malleolar markers would have the advantage, compared to using the lateral malleolar or the fifth metatarsal marker, of considering the orientation of the foot (Tisserand et al., 2018, 2016), especially in individuals with consequent internal or external foot rotation (example in **figure 1**). Regardless of the approach chosen, the marker representing the BoS edge is often placed on a foot edge that is not always in contact with the ground (e.g., the lateral malleolar marker during late stance), which limits its effectiveness because only a body part in contact with the ground can contribute to quickly move the CoP. To overcome this problem, the marker defining the edge of the BoS could be chosen according to two conditions: **1)** the marker should be the most anterior (for the AP edge of the BoS) or the most lateral (for the ML edge of the BoS), and **2)** be fixed on a physical edge of the foot that is in contact with the ground at the instant when the MoS is calculated. To date, no study has investigated the effect of employing these various approaches of defining the edges of the BoS, which could potentially induce significant differences in the MoS measure, particularly for the ML MoS.

**Figure 1.**
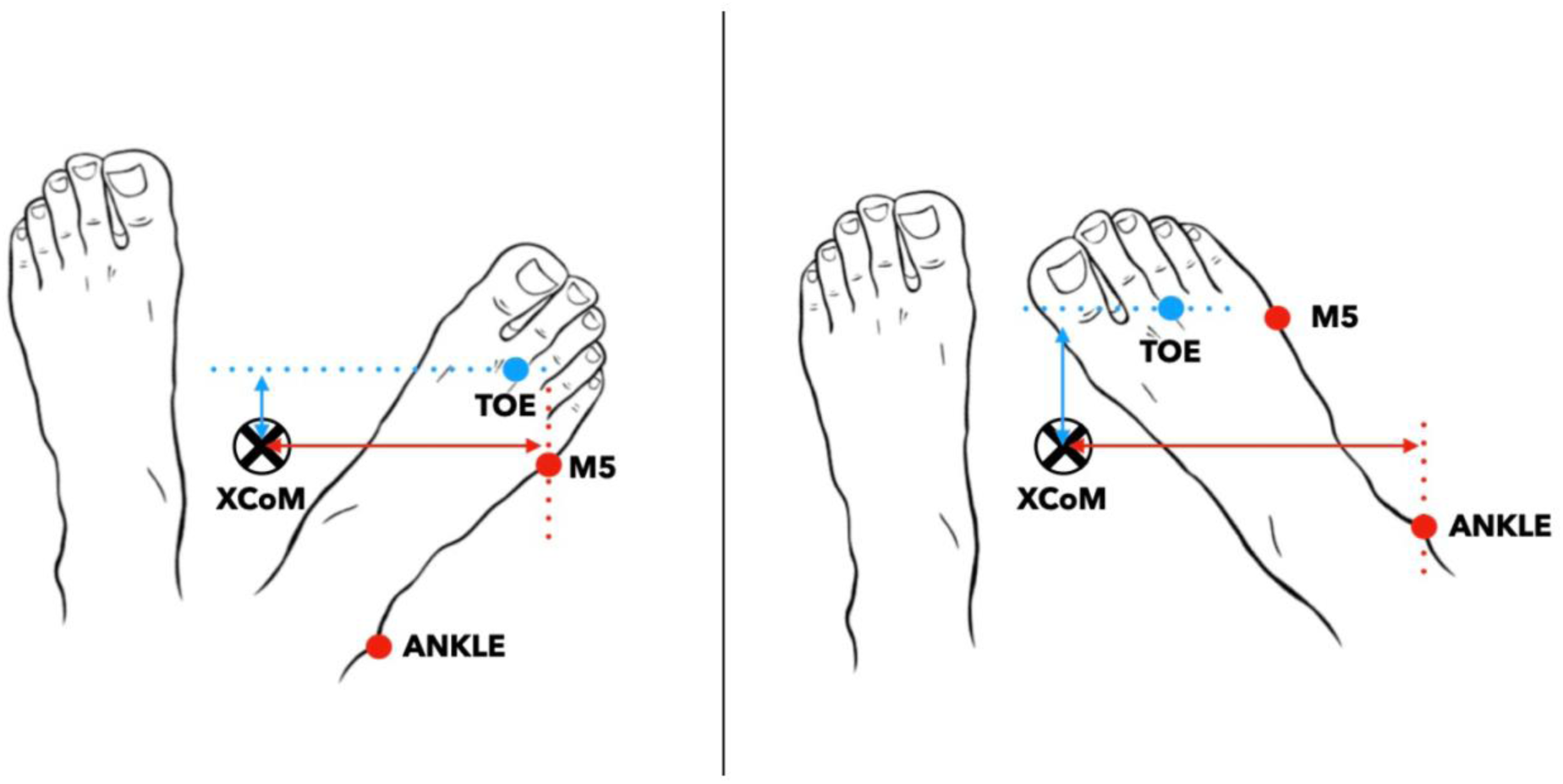
Representation of the most lateral marker during foot rotation between the fifth metatarsal (M5) and lateral malleolar (ANKLE) markers. The medio-lateral margin of stability is the difference between the extrapolated center of mass (xCOM) and the most lateral marker. The antero-posterior margin of stability is the difference between xCOM and the most anterior marker, which is always the second metatarsal head marker (TOE). The medio-lateral margin of stability is the difference between the extrapolated center of mass (xCOM) and the most lateral marker.

In addition to being calculated heterogeneously (mostly in the ML direction), the MoS is typically computed at a single time point, specifically at the initial contact of the ipsilateral foot (Watson et al., 2021). Calculating the MoS at this point is crucial as it reflects the mechanical effects of the contralateral stance phase. However, it has been observed that the minimal MoS is achieved before the contralateral toe-off (Hof, 2007; Hof et al., 2005), and later suggested to be the instant when the MoS should be measured (Curtze et al., 2024).

The linear model used to calculate the MoS includes few limitations, which arise from simplifying assumptions made to facilitate biomechanical analysis such as the unchanging pendulum length, the non-deformable pendulum, and the non-consideration of potential additional external forces. Since the MoS accounts for the CoM velocity in its calculation, the AP MoS is strongly influenced by gait speed, with more negative AP MoS when gait speed increases (Curtze et al., 2024; McCrum et al., 2019). Indeed, the AP MoS remains negative during the stance phase of a steady-speed gait because the BoS is consistently positioned posterior to the xCoM (Curtze et al., 2024; Hof, 2008). Mainly, the body is always “falling forward,” and stability is maintained by continuous repositioning of the BoS, as the foot contacts the ground in the next step (Kuo and Donelan, 2010). However, the influence of gait speed on ML MoS is less understood and has been investigated by fewer studies (Gates et al., 2013; Guaitolini et al., 2019; Peebles et al., 2016). Moreover, the MoS model neglects the influence of anthropometric factors, such as foot length, foot width, and body mass. Since the MoS is a widely used clinical measure of gait stability and is often compared across populations with different anthropometric characteristics and gait speed, the effect of these factors should be investigated to better personalize/individualize clinical assessments.

This study aimed to 1) assess differences between AP and ML MoS measures resulting from the different approaches of defining the edges of the BoS that have been previously used in the literature and the one proposed in this current study, and 2) investigate the effect of foot length, foot width, gait speed, and body mass on the MoS measures. It was hypothesized that 1) the different approaches of defining the BoS used will lead to different AP and ML MoS values, and that 2) AP MoS measure will be negatively related to foot length, gait speed, and body mass, and ML MoS will be positively related to foot width, gait speed, and body mass.

## 1. Method

### 1.1. Participants

This study used an open access dataset from Riglet et al. (2024)’ study, including 30 healthy participants (16M/14F) aged between 21 and 41 years old (27.97 ± 5.59 years old) (Riglet et al., 2024). Participants were on average 1.73 ± 0.92 m tall and weighed 68.16 ± 11.06 kg.

### 1.2. Procedure

#### Records

Each participant was instructed to walk at three different speeds along 10 meters of a flat and regular laboratory surface: slow, comfortable, and fast with walking shoes provided for the experiment (Riglet et al., 2024) (**supplementary figure 1**). All participants were equipped with a set of 63 reflective markers following the Conventional Gait Model (pyCGM, v.2.5) (Baker et al., 2018). Eighteen optoelectronic cameras (Vicon System®, Oxford, UK; 100 Hz sampling rate) and Nexus software were used to collect the marker trajectories.

#### Data processing

The c3d files that include the labelling of the marker trajectories were then exported from the Riglet et al. (2024) (Riglet et al., 2024) database and further processed in MATLAB (vR2022b, Mathworks Inc., Natick, USA) using the open-source biomechZoo toolbox (v.1.9.10) (Dixon et al., 2017) and custom codes. The foot strikes and foot-offs were identified using the method of Zeni et al. (2008), based on the positional changes of the heel, foot, and sacrum markers (Zeni et al., 2008). Then, the walking trials were partitioned into individual gait cycles. Considering the natural asymmetry in able-bodied gait (Sadeghi et al., 2000), gait cycles of both legs were included. For each participant, the first 12 gait cycles of each walking speed condition were used for the MoS calculation (i.e., 6 gait cycles on each leg).

#### Calculations

For each gait cycle, the continuous MoS was calculated during the stance phase in the AP and ML directions using **Eq.1** and **Eq. 2** (Hof et al., 2005). The stance phase started at the ipsilateral foot contact and ended at the ipsilateral foot-off. The anterior direction of walking was described as the vector of the walking direction in the transverse plane of the laboratory whereas the lateral direction of walking was described as the vector perpendicular to the anterior direction of walking. A table summarizing the different approaches used in the previous literature to calculate the AP and ML MoS is provided in the appendix (**supplementary table 1**).

The AP MoS was calculated at each time point of the stance phase following 3 approaches that have been used in the previous literature (HEEL, TOE, MOST POSTERIOR):

1. **HEEL:** Using the heel marker (**HEEL**) as the anterior limit of the BoS.
2. **TOE:** Using the second metatarsal marker (**TOE**) as the anterior limit of the BoS.
3. **MOST ANTERIOR:** Using the most **anterior** marker between the **HEEL** and **TOE** markers. The part of the foot to which the most anterior marker is attached had to be in contact with the ground. For instance, at foot strike, the **HEEL** marker was chosen if the forefoot (**TOE**) was elevated, whereas the **TOE** marker was selected if the heel was elevated (i.e. during late stance).

The ML MoS was calculated at each time point of the stance phase following 4 approaches (ANKLE, M5, MIDPOINT, MOST LATERAL):

1. **ANKLE:** Using the lateral malleolar marker (ANKLE) as the lateral limit of the BoS.
2. **M5:** Using the fifth metatarsal marker (**M5**) as the lateral limit of the BoS. By identifying the most lateral marker as the lateral limit of the BoS.
3. **MIDPOINT:** Using the midpoint of the virtual line relating the **ANKLE** and **M5** markers as the lateral limit of the BoS.
4. **MOST LATERAL:** Using the most **lateral** marker between the **ANKLE** and **M5** markers. The part of the foot to which the most lateral marker is attached had to be in contact with the ground. For instance, at foot strike, the **ANKLE** marker was chosen if the midfoot/forefoot (**M5**) was elevated, whereas the **M5** marker was selected if the heel was elevated (i.e. during late stance).

All calculations were performed using MATLAB (vR2024a, Mathworks Inc., Natick, USA). For each gait speed condition, calculation approach, and participant, the MoS curves were averaged.

Gait speed was calculated as the distance covered by the head marker during a gait cycle divided by the gait cycle duration in seconds. The foot length was provided within the Riglet et al. (2024) database (Riglet et al., 2024), and the foot width was calculated as the distance between the first and the fifth metatarsal markers.

### 1.3. Statistical analysis

All statistical analyses in this study were carried out using the Statistical Parametric Mapping (SPM) toolbox (Pataky, 2010) and using custom-made Matlab scripts.

To test our first hypothesis, a paired t-test was conducted to assess differences between the of the AP MoS calculation approaches (i.e., HEEL and TOE). To evaluate the differences between the three ML calculation approaches (i.e., ANKLE, M5, and ANKLE-M5), a repeated-measures Analysis of Variance (ANOVA) was performed. In the presence of significant results for the latter, post-hoc analyses using paired t-test were carried out. The level of significance was adjusted following a Bonferroni correction (n = 3) to account for multiple comparisons (i.e. M5 vs MIDPOINT, M5 vs ANKLE, ANKLE vs MIDPOINT) (Altman, 1990). The mean Cohen’s d (*d*) effect size was calculated for each significant cluster (i.e., intervals during the stance phase when the p-value is below the threshold) (Cohen, 1977). Only clusters lasting 5% of the stance phase or more were discussed (Armijo-Olivo et al., 2011). Only the MoS curves at comfortable speed were included (n = 30 curves).

To test our second hypothesis, a correlation analysis was conducted to assess the relation of foot length, gait speed, and body mass with the continuous AP MoS curves using the SPM toolbox (spm1d.stats.regress function, spm1d version M.0.4.11). The same analysis was conducted to assess the relation of foot width, gait speed, and body mass with the continuous the ML MoS curves. The AP MoS and the ML MoS were calculated using the MOST ANTERIOR and the MOST LATERAL approaches, respectively. For all the significant clusters, the mean coefficient of correlation (*r*) was calculated. For the gait speed correlation analysis, the MoS curves of each condition (i.e., slow, comfortable, fast) were included (n = 90 curves). The relationship was interpreted as small (*r* = 0 – 0.290), moderate (*r* = 0.300 – 0.490), strong (*r* = 0.500 – 0.69), and very strong (*r* = 0.700 – 1.000). Differences in gait speed for each condition are presented in **supplementary figure 1**.

## 2. Results

### 2.1. Differences between the calculation approaches

The calculation approach yielded to different results throughout the entire stance phase: HEEL vs TOE (0 – 100%, p < 0.001, *d* = −1.297) (**figure 2a, 2b**). When using the MOST ANTERIOR approach, the HEEL marker was used at the beginning of the stance phase (0 – 20%) after which the TOE marker was selected (**figure 2a**).

**Figure 2.**
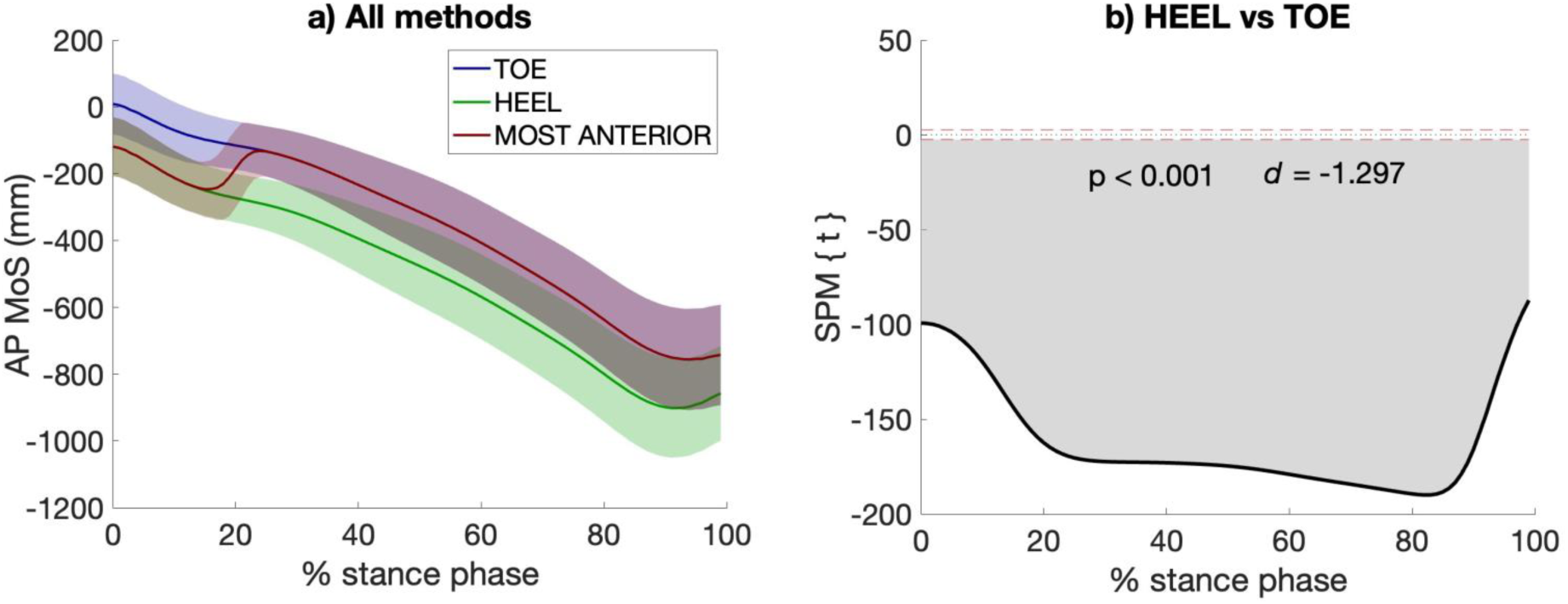
Antero-posterior (AP) margin of stability (MoS) calculated using the two most widely used approaches in the literature (i.e., TOE, HEEL), as well as the approach proposed in this study (i.e., MOST ANTERIOR) to describe the anterior limit of the base of support (**a**). A negative AP MoS refers to an extrapolated center of mass that is in front of the anterior limit of the BoS. The differences between approaches are presented (statistical parametric mapping (SPM) paired t-test, p < 0.05) (**b**). The red dashed lines indicate the critical thresholds for statistical significance. Values above or under these lines indicate statistically significant differences between the compared approaches at that specific point in the stance phase.

Concerning the ML MoS, the ANOVA showed a significant difference between all calculation approaches (p < 0.001) over the entire stance phase (**supplementary figure 2**). Consequently, paired t-tests were performed. Compared to each other, all calculation approaches lead to different ML MoS values throughout the stance phase: ANKLE vs M5 (0 – 100%, p < 0.001, *d* = 1.701), ANKLE vs MIDPOINT (0 – 100%, p < 0.001, *d* = 0.887), M5 vs MIDPOINT (0 – 100%, p < 0.001, *d* = −0.866) (**figure 3b, 2c, 2d**). When using the approach of the most lateral marker, the ankle marker was chosen during the first 20% of stance phase. Else, the fifth metatarsal marker was chosen as the most lateral marker (**figure 3a**).

**Figure 3.**
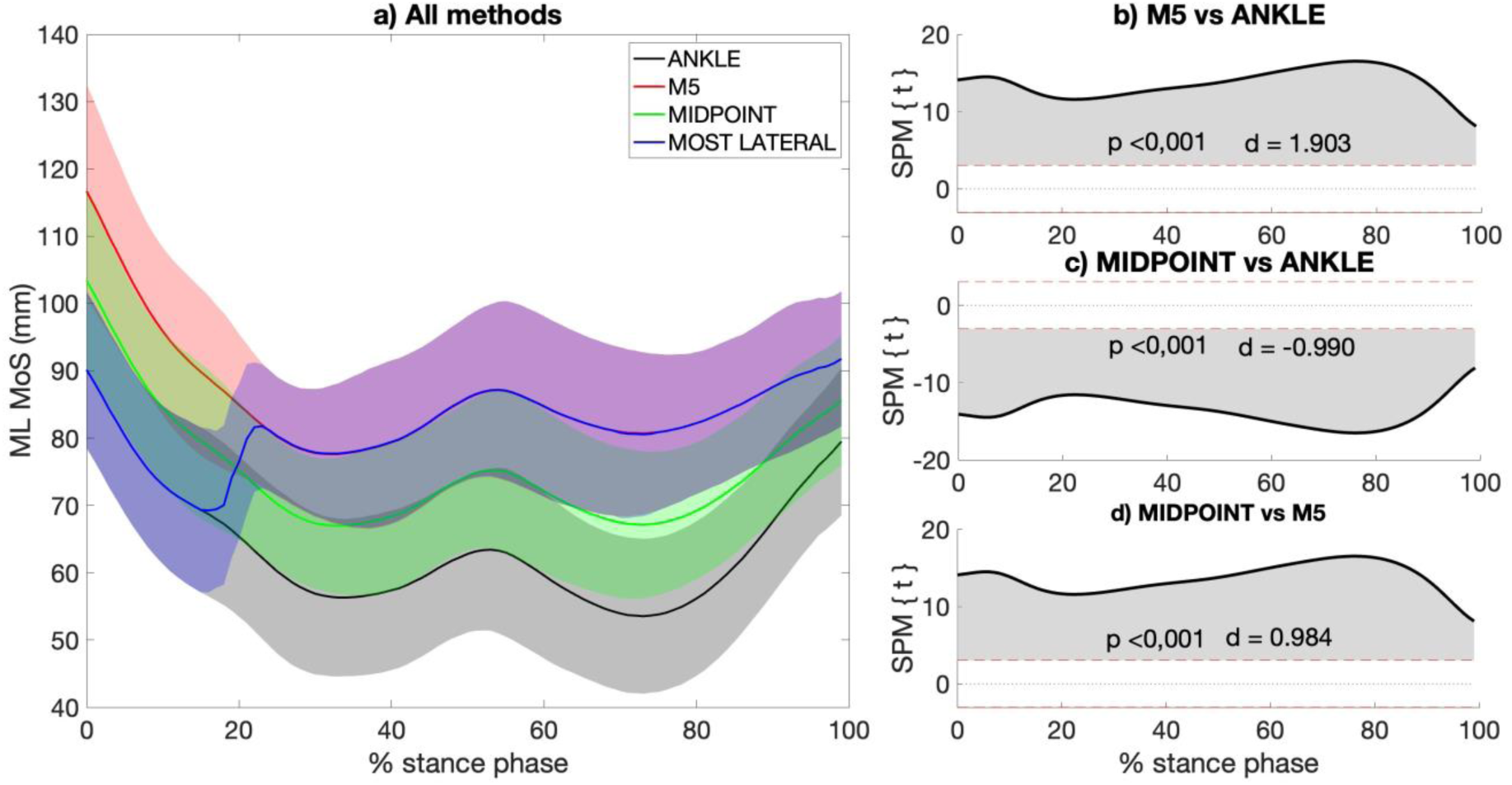
Mediolateral (ML) margin of stability (MoS) calculated using the three most widely used approaches in the literature (i.e., ANKLE, M5, MIDPOINT), as well as the approach proposed in this study, (i.e., MOST LATERAL) to describe the lateral limit of the base of support **(a)**. A negative ML MoS refers to an extrapolated center of mass that is more lateral than the lateral limit of the BoS. The differences between approaches are presented (statistical parametric mapping (SPM) paired t-test, p < 0.05) **(b, c, d)**. The red dashed lines indicate the critical thresholds for statistical significance. Values above or under these lines indicate statistically significant differences between the compared approaches at that specific point in the stance phase.

### 2.2. Factors influencing the margin of stability

The AP MoS was negatively correlated with foot length (8 – 18% of stance phase, p = 0.029, *r* = −0.559; 23 – 100%, p < 0.001, *r* = 0.640), gait speed (0 – 100%, p < 0.001, *r* = −0.905), and body mass (11 – 18%, p = 0.038, *r* = −0.524; 24 – 100%, p < 0.001, *r* = −0.598) (**figure 4a, 4c, 4e**). The ML MoS was positively correlated with gait speed (22 – 30%, p = 0.033, *r* = 0.449; 78 – 100%, p = 0.003, *r* = 0.411), body mass (66 – 97%, p < 0.001, *r* = 0.563), but not foot width (**figure 4b, 4d, 4f**). The detailed results of the correlation analysis are presented in the **supplementary figure 3 and 4**.

**Figure 4.**
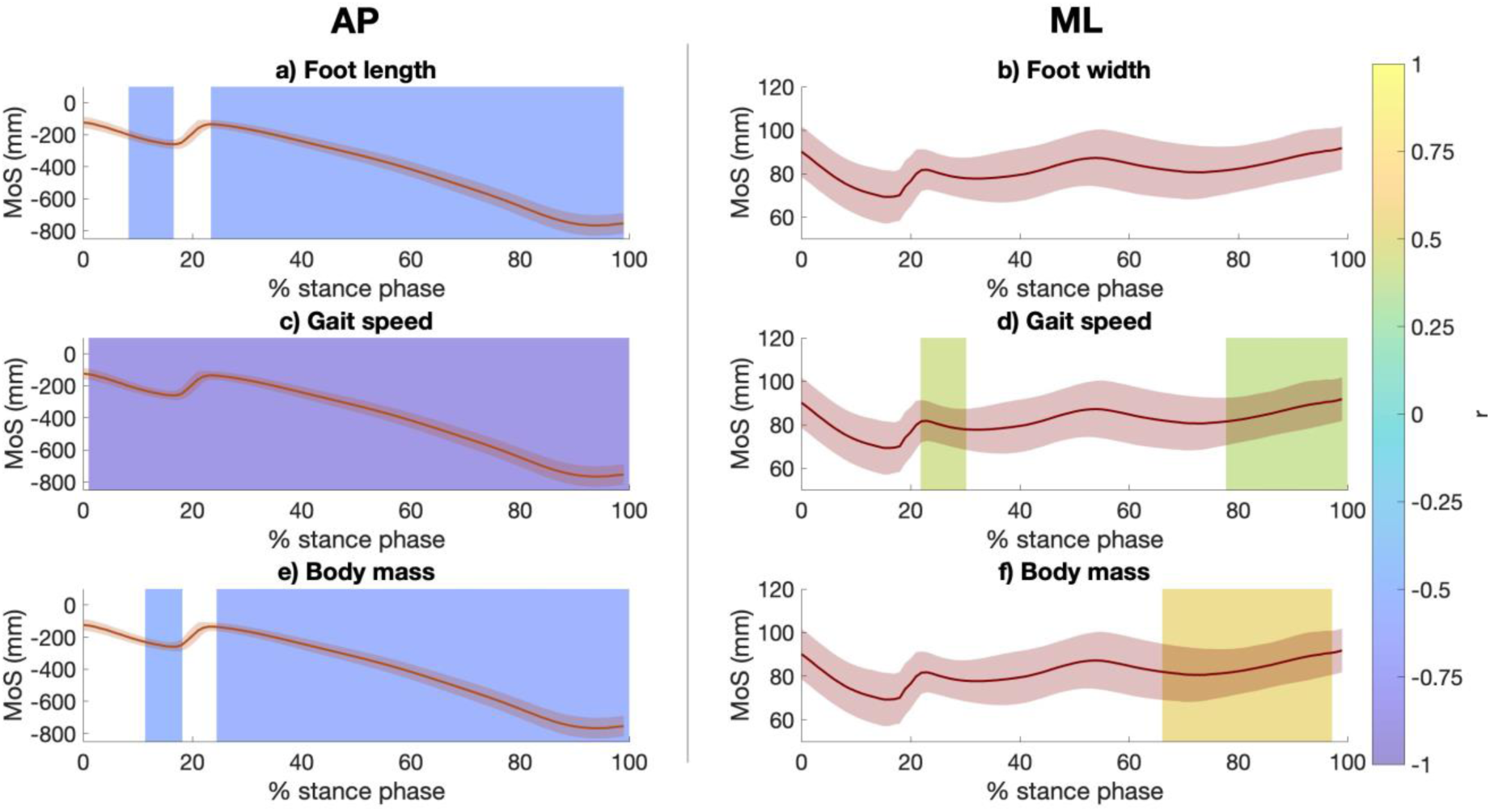
Correlation analysis results between foot length **(a)** or width **(b)**, gait speed **(c, d)**, and body mass **(e, f)** and the margin of stability (MoS) in the anteroposterior (AP) **(a, c, e)** and mediolateral (ML) **(b, d, f)** directions during the stance phase of the gait cycle. The AP MoS was calculated using the MOST ANTERIOR approach whereas the ML MoS was calculated using the MOST LATERAL approach. A negative AP MoS refers to an extrapolated center of mass that is in front of the anterior limit of the BoS, whereas a negative ML MoS refers to an extrapolated center of mass that is more lateral than the lateral limit of the BoS. The colored patches represent regions where significant correlation were found (statistical parametric mapping, p < 0.05), and are colored by its mean *r* value.

## 3. Discussion

### 3.1. Summary

The aims of the present study were to 1) assess differences between the approaches most widely used in the literature for calculating AP and ML MoS and 2) investigate the effect of foot length, foot width, gait speed, and body mass on the MoS measures. Two main findings immerged. First, significant differences were observed between the MoS calculation approaches (using the HEEL and TOE markers for AP MoS, and using the M5, ANKLE, and MIDPOINT for ML MoS). Second, gait speed, foot length, and body mass are negatively correlated with the AP MoS during almost the entire stance phase, whereas only gait speed and body mass are positively correlated with the ML MoS and only during late stance.

### 3.2. The effect of the calculation approach

A key component for calculating the MoS is the BoS. The latter can be defined as the area between the feet during walking outlined by the points of contact with the ground. Placing a marker on specific points such as medial or lateral malleolus, M5, or calcaneus might position the BoS edge too far toward the center, sides, front, or back. Consequently, this can result in misestimations of the ML and AP MoS (Curtze et al., 2024). The results of the present study showed that the marker chosen to describe the anterior or lateral BoS boundaries significantly affects the resulting values of AP and ML MoS, respectively (**figure 2** and **figure 3**). Our results complement those of Havens et al. (2018), who have reported that biases in the MoS value can be introduced by the approach used to estimate the CoM dynamics (Havens et al., 2018). The current study used the pelvis average model (i.e., average position of the four pelvis markers) to minimize this bias, as suggested by the latter study (Havens et al., 2018) when simplified models of CoM dynamics are used. Together, these results highlight that the MoS can be significantly affected by the choice of calculation approach, both in terms of the definition of the BoS and the method used to estimate the CoM kinematics. This underscores the importance of avoiding comparisons between studies that use different calculation approaches and the future adoption of a standardized approach across different studies.

The current literature reveals an important heterogeneity in MoS calculation, which makes comparisons between studies and populations difficult (Watson et al., 2021). For example, in children with cerebral palsy, some studies opted for the lateral malleolus to describe the lateral boundary of the BoS (Delabastita et al., 2016; Ma et al., 2021; Rethwilm et al., 2021), while other used the MIDPOINT approach (Sangeux et al., 2024). In populations with in-toeing or out-toeing gait such as those with cerebral palsy (Cao et al., 2020; Rethlefsen, 2006), both approaches may lead to misestimates the BoS (Puszczałowska-Lizis and Ciosek, 2017). Similarly, using the M5 marker to describe the lateral boundary of the BoS may not be relevant if the individual presents a rotation of the medial foot. Thus, these approaches to calculate the MoS are less suited to pathological populations, especially when the MoS value is used to be compared with their healthy peers. Our study reported a significant difference in the AP and ML MoS values based on the choice of the markers used to define the BoS in non-pathological gait. Finally, another potential limitation of the current literature is the calculation of the MoS at a single point in the gait cycle, often at initial contact (Hak et al., 2013b; Rijken et al., 2015; Sangeux et al., 2024), although it has been largely suggested focus on more relevant instant, such as close to the contralateral toe-off (Curtze et al., 2024), which is when the MoS is minimal and stability is mechanically critical (Hof, 2007; Hof et al., 2005).

Treating MoS as a continuous measure has been emphasized in the literature, such as McAndrew Young et al. (2012) who highlighted the importance of studying the MoS throughout the entire stance phase instead of an average value across the gait cycle, to better highlight the instant when the stability is critical (McAndrew Young et al., 2012). Similarly, Kazanksi et al. (2022) proposed a step-to-step approach to solve the MoS averaging paradox (Kazanski et al., 2022). These findings support our goal of clarifying MoS interpretation.

### 3.3. The effect of foot length and width

The findings of this study indicate that the MoS is more forward in individuals with longer feet (**figure 4a**), which may reflect an adaptation to the individual’s longer limb structure. This correlation has been noted even before the foot achieves full contact (i.e., between 8 – 18 % of stance phase), suggesting that foot morphology such as foot length may play a role in dynamic stability even when the foot is not completely in contact with the ground. Indeed, with a longer foot, the CoP may extend further from the ankle compared to individuals with smaller feet, potentially allowing for greater ankle moment to help regulate the stability, and in cases of significant instability, to decelerate. To our knowledge, this is the first study that have assessed the effect of foot length on AP MoS during gait (**supplementary table 1**), which limits our understanding on how a forward MoS may promotes a more stable gait in individuals with longer feet. However, as the MoS is a measure of an individual gait stability and is commonly compared between populations, normalizing this value by foot length should be considered to ensure accurate comparisons.

Regarding the ML direction, it was expected that larger foot will lead to wider BoS, which could have directly increased the ML MoS. However, no significant relation has been noted between foot width and the MoS (**figure 4b**). During postural balance task (i.e., eyes open and closed), Qiu et al. (2013) observed that wider foot width is related to greater balance performance (measured by the Composite Equilibrium Score), especially in younger individuals (Qiu and Xiong, 2013). Together, these results may suggest that the mechanics of gait stability, in contrast to postural balance tasks, involve more complex factors beyond just foot width especially in the ML direction (Bauby and Kuo, 2000; Kuo and Donelan, 2010).

### 3.4. The effect of gait speed

It has been also observed that individuals used a more forward MoS when gait speed is increased, which was characterized by a very strong correlation coefficient (*r* = −0.887) (**figure 4c**). It is important to recognize that this strong correlation is likely influenced by the fact that the calculation of xCoM incorporates a velocity component. Nevertheless, other previous studies have also indicated that gait speed plays a key role in an individual’s AP stability (Guaitolini et al., 2019; Hak et al., 2013a; McCrum et al., 2019). Indeed, consistent with our findings, Guaitolini et al. (2018) reported that an increase in the gait speed is related to a forward MoS (i.e., the xCoM is more forward relative to the BoS) in healthy young adults (Guaitolini et al., 2019). Thus, normalization of the MoS by gait speed should be considered to reduce between-participant variability, which is an approach that has been advocated previously to enhance the precision of stability assessments (McCrum et al., 2019).

In the ML direction, an increased gait speed was moderately related to a larger MoS (i.e., the xCoM is more medial than the lateral limit of the BoS), during mid-stance (22 – 30%, *r* = 0.449) and late stance (78 – 100%, *r* = 0.411) (**figure 4d**). Especially during late stance, this result can be explained by the reduced ML oscillation amplitude of the CoM when walking faster, as compared to slower walking speeds. This reduced oscillation would result in a lower ML velocity, which in turn increases the ML MoS. However, this finding was not supported by Lencioni et al. (2019), who reported a negative correlation (*r* = −0.320) between gait speed normalized by body height and the ML MoS at mid-stance in healthy adults (Lencioni et al., 2020). The moderate relationship suggests that other factors such as stance time and step width may play a more significant role in influencing the ML MoS during mid-stance and late stance (Buurke et al., 2023; Hof, 2008). Also, the weaker relationship (i.e., lower correlation coefficient) in the ML compared to the AP direction, aligns with the principle that the xCoM incorporates the velocity of the CoM. Consequently, the AP MoS is more strongly related to gait speed than the ML MoS since more motion occurs in that direction (McCrum et al., 2019). A previous study has shown that the ML MoS may be, among other factors, related to age (i.e., older individuals will have a larger MoS) (Lencioni et al., 2020). However, the present study did not assess the relationship between age and ML MoS, due to the narrow range of age (21-41 years old) of the included population.

### 3.5. The effect of body mass

A more forward MoS was observed when body mass is increased during the end of the initial contact phase (11 – 18%, *r* = −0.523), and during foot flat and push-off phases (25 – 100%, *r* = −0.598) (**figure 4e**). This observation may align with previous finding that obese individuals tend to adopt a more forward posture during balance-challenging tasks, which has been attributed to the increased difficulty they encounter in controlling anterior-posterior body movements (Berrigan et al., 2006; Menegoni et al., 2009). In the ML direction, a larger MoS (i.e., the xCoM is more medial than the lateral limit of the BoS) was observed in individuals with increased body mass during late stance (66 – 97%, *r* = 0.563) (**figure 4f**). This may reflect a strategy for conserving energy, possibly driven by their larger mass, which reduces the need for substantial mechanical effort to shift to the other side. This adaptation aligns with the findings of previous studies who also reported a larger MoS during gait in healthy individuals with higher body mass (Herssens et al., 2020; Lencioni et al., 2020).

### 3.6. Limitations

This study has some limitations. First, the MoS is an instant measure or gait stability that does not allow for a comprehensive understanding of the factors contributing to gait stability such as segmental/joint kinematics and muscle activity. Future research should consider integrating these parameters for a more refined interpretation of the MoS. Second, expanding the study to include pathological populations and a wider age range would help generalize the findings and enhance their clinical relevance. Our approach is intended to be inclusive of pathological populations, but we have not yet validated it with these groups. Third, the age factor is also underrepresented, as the participant age range is too narrow, limiting the applicability of the results to other age groups.

## 4. Conclusion

This study stands out as one of the few to have continuously evaluated the MoS across the gait cycle. It introduces a novel method using optimal foot markers to estimate BoS in the absence of CoP data. By consistently normalizing MoS (AP by foot length, gait speed, and body mass; ML by gait speed and body mass), this approach enhances gait stability assessment, improves population comparisons, and better identifies stability deviations in pathological gait patterns.

## Author contributions

Conceptualization; CDP, YC - Formal analysis; CDP, CR, YC - Methodology; CDP, CR, YC - Supervision; YC - Visualization; CDP, CR, YC - Roles/Writing - original draft; CDP, YC - Writing - review & editing; YC, RT

## Acknowledgements

The authors would like to acknowledge the Fonds de recherche du Québec (FRQ)—Nature et technologie, for the doctoral funds of the first author.

---

**Supplementary figure 1.**
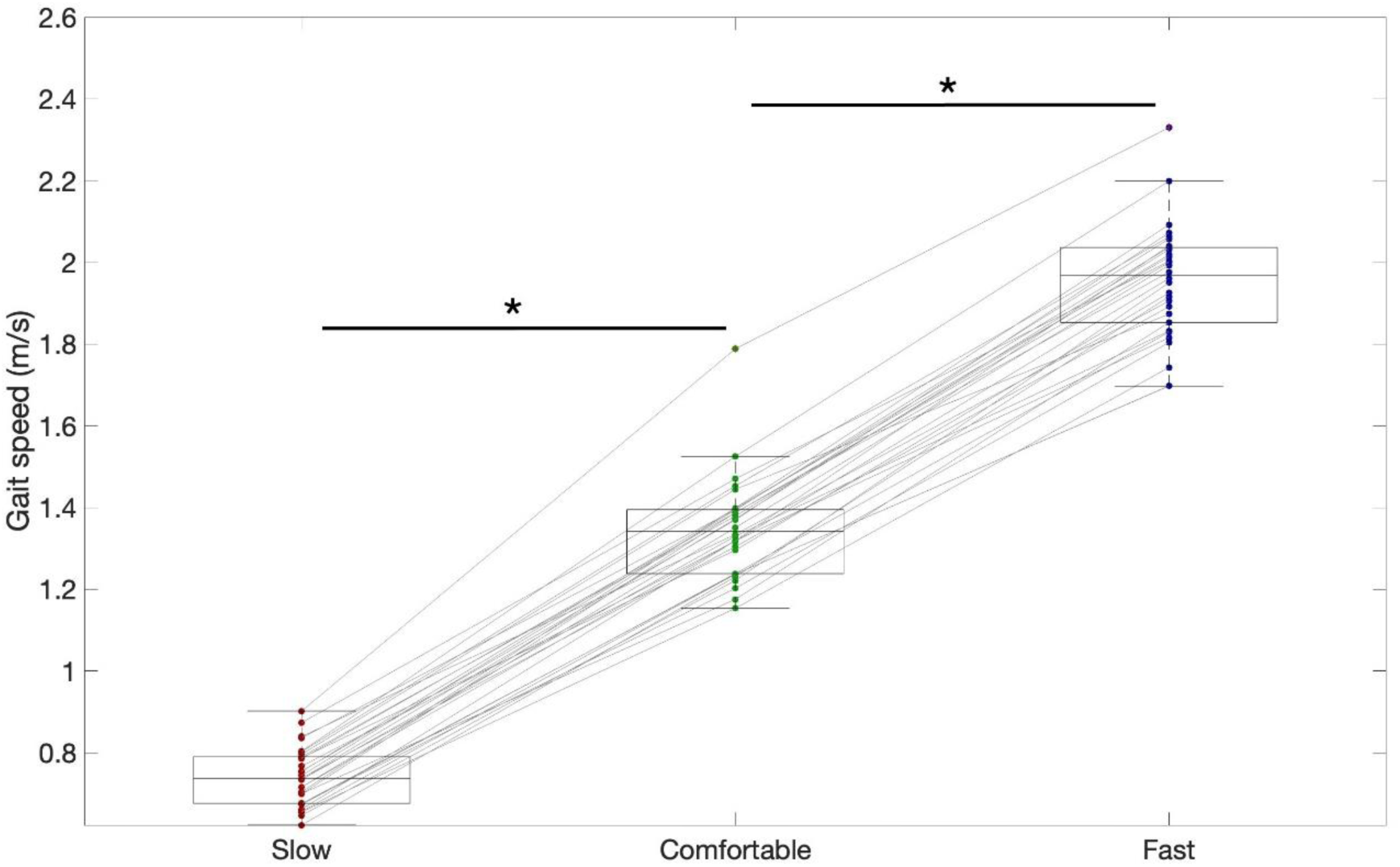
Boxplots representing the participant mean gait speed (m/s) across 12 gait cycles for each condition (slow, comfortable, fast), with the red line indicating the group median gait speed. Individual changes across the three speed conditions are presented with a grey line that connects the mean gait speed. The top and bottom edges of the box represent the first quartile (Q1) and third quartile (Q3), respectively. The whiskers extend to the largest and smallest values within 1.5 times the interquartile range from Q3 and Q1, respectively. The black horizontal lines with asterisks (*) denote statistically significant differences (p < 0.05, Wilcoxon Signed-Rank test) between the speed conditions. Participants gait speed were significantly different between speed conditions: slow vs comfortable (median difference [95% confidence interval (CI)]: 0.60 [0.56, 0.64] m/s, p < 0.001, Glass delta effect size (△): 4.930); comfortable vs fast (median difference [95% CI]: −0.61 [-0.66, −0.59] m/s, p < 0.001, △: 4.934) (see figure 2).

**Supplementary figure 2.**
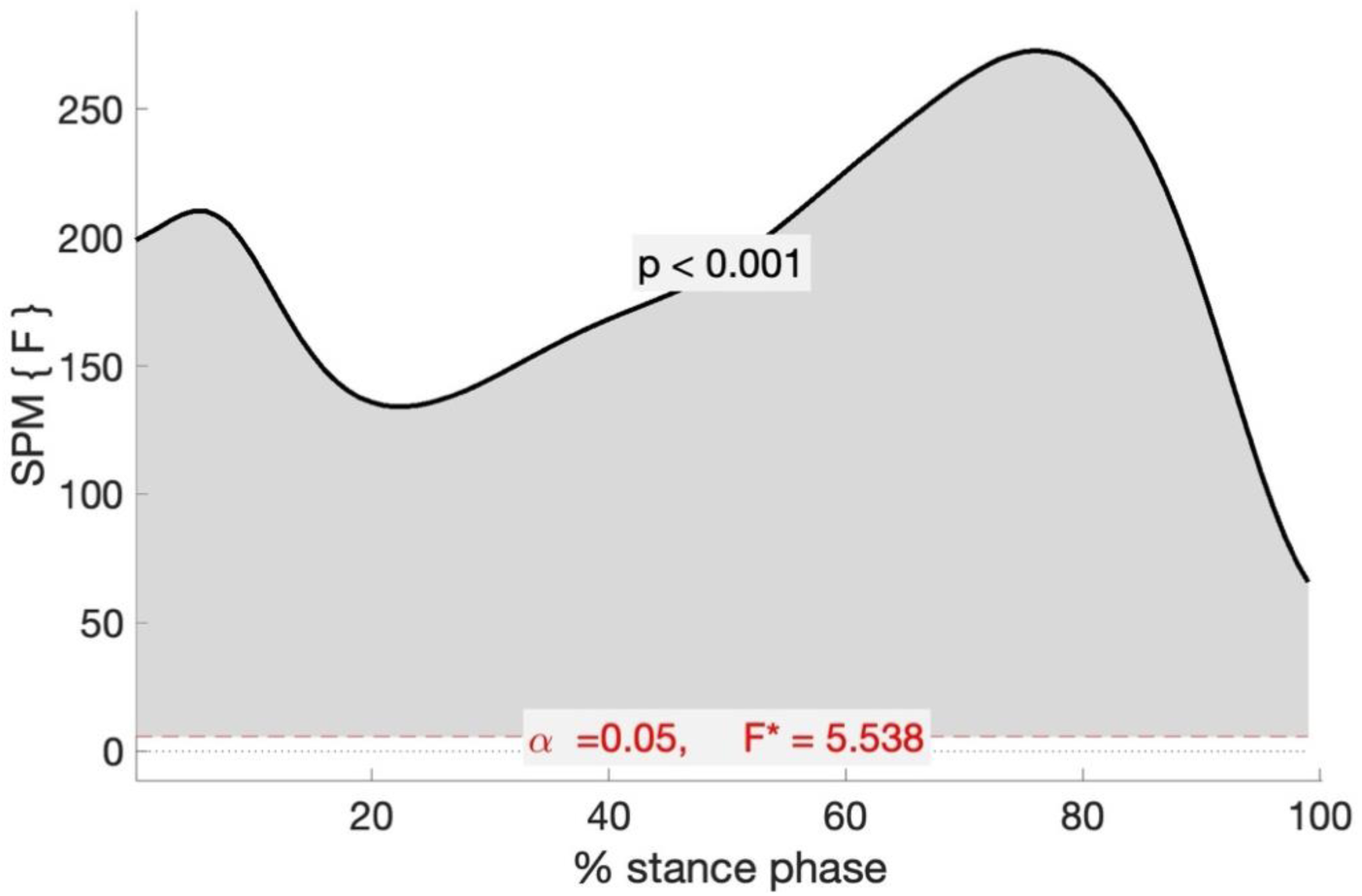
Results of the statistical parametric mapping analyses investing the analysis of variance (ANOVA) between the mediolateral margin of stability calculation approaches. F-statistic from the ANOVA, used to determine whether there are significant differences between the three groups is represented by the red dashed lines as the critical thresholds for statistical significance. Values above or under these lines indicate statistically significant difference (ANOVA, p < 0.05) between the compared approaches at that specific point in the stance phase.

**Supplementary figure 3.**
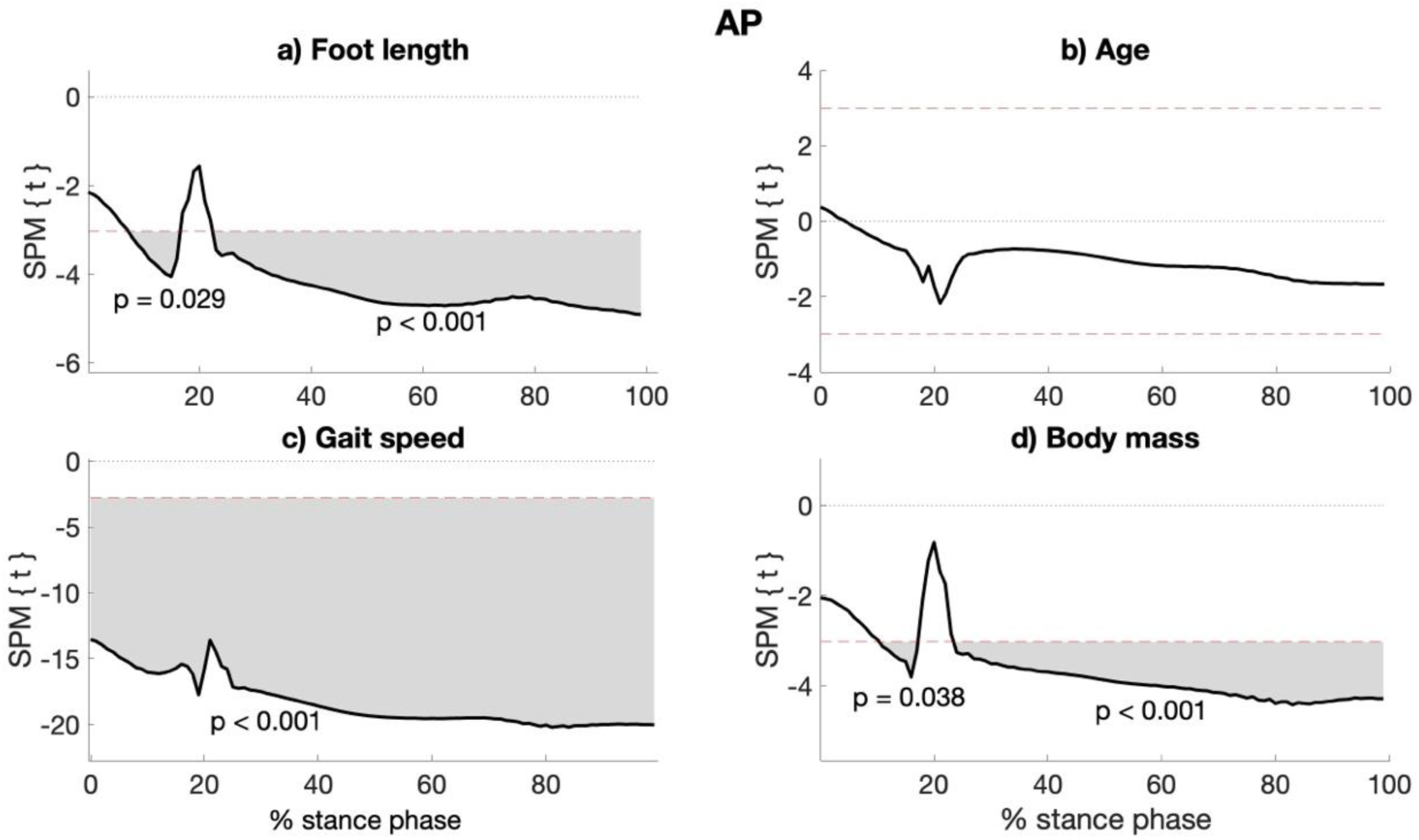
Results of the statistical parametric mapping (SPM) analyses investigating the regression between foot length (a), age (b), gait speed (c), and weight (d), and the antero-posterior (AP) margin of stability throughout the stance phase of the gait cycle.

**Supplementary figure 4.**
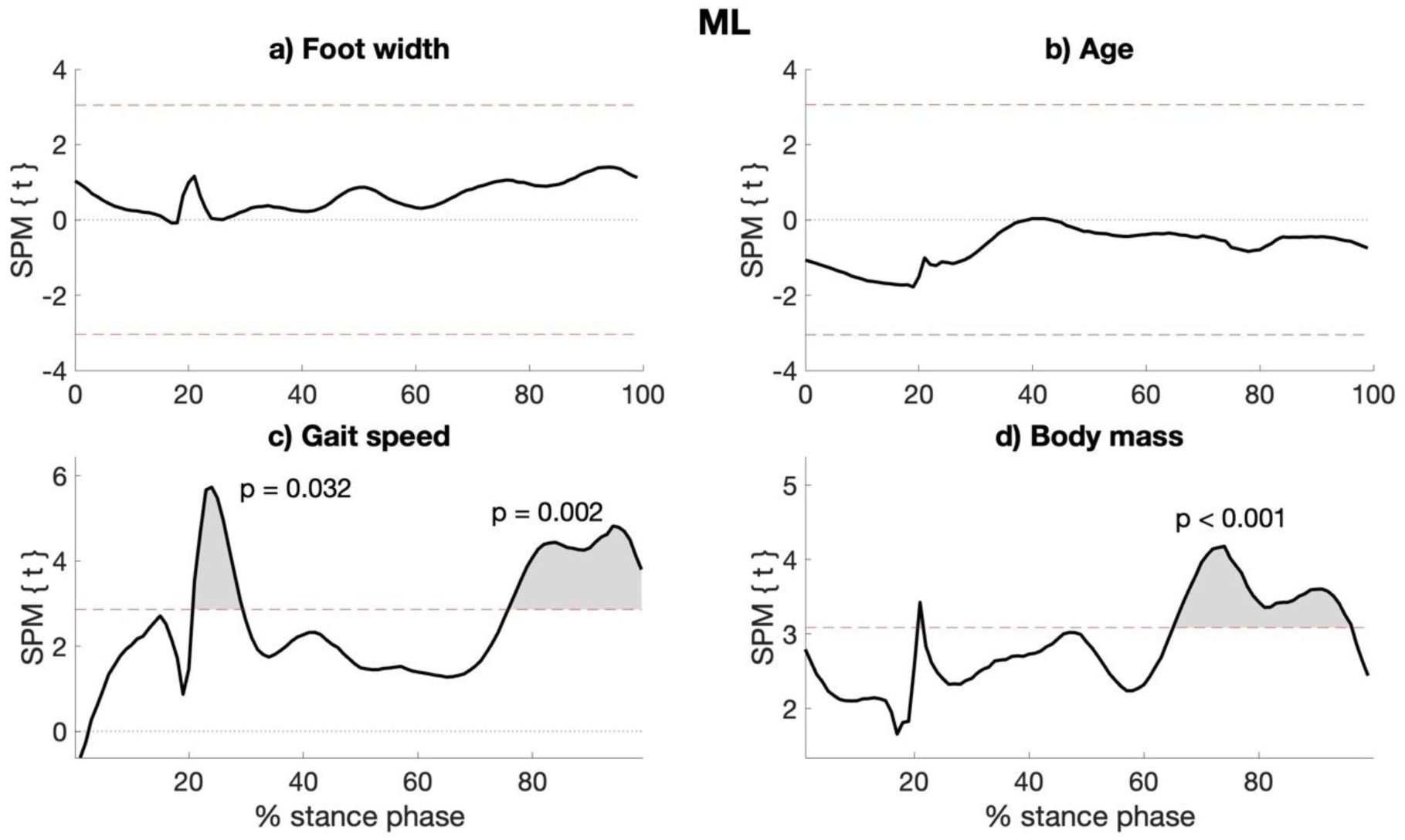
Results of the statistical parametric mapping (SPM) analyses investigating the regression between foot length (a), age (b), gait speed (c), and weight (d), and the medio-lateral margin (ML) of stability throughout the stance phase of the gait cycle.

**Supplementary table 1.**
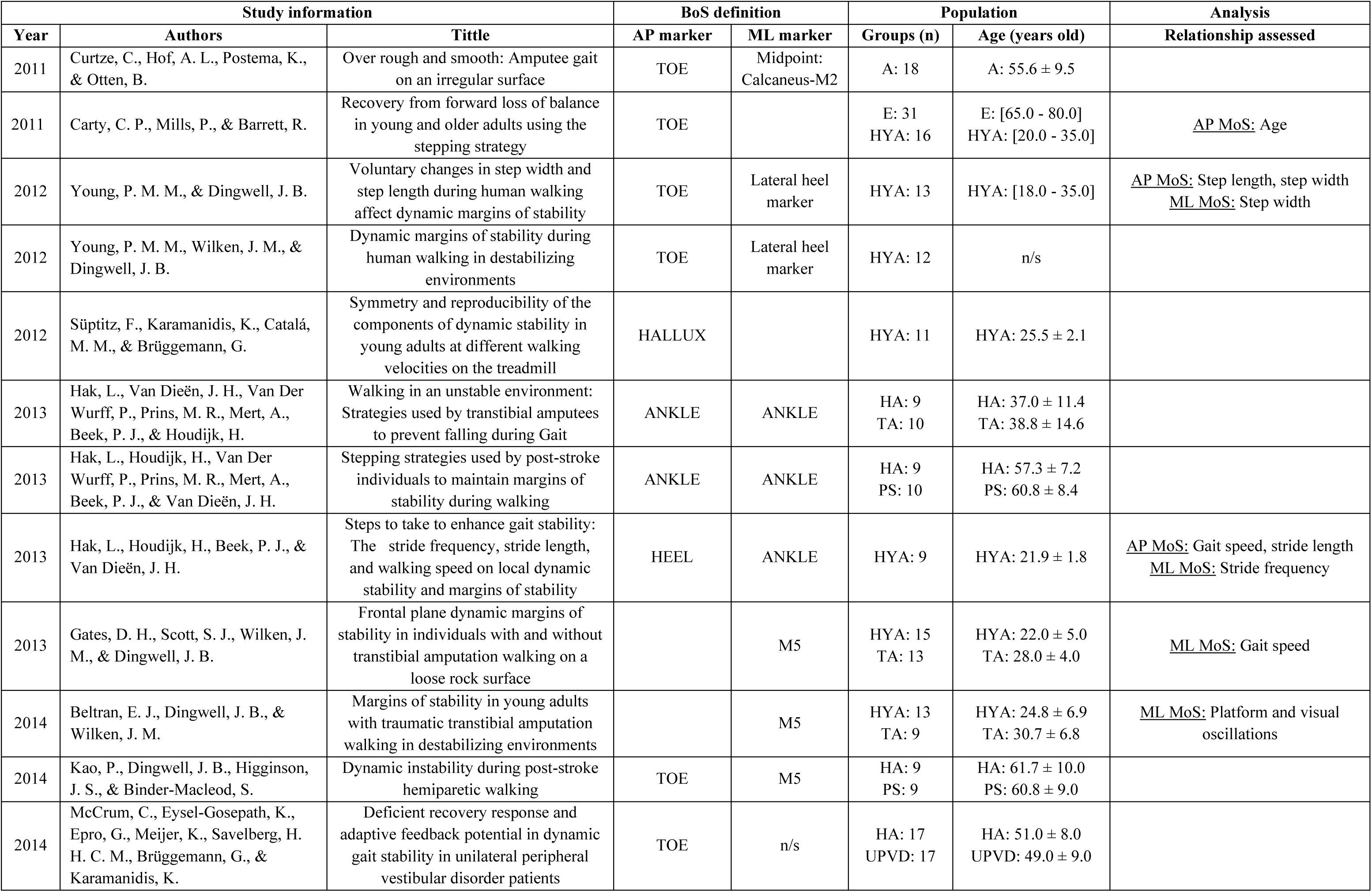
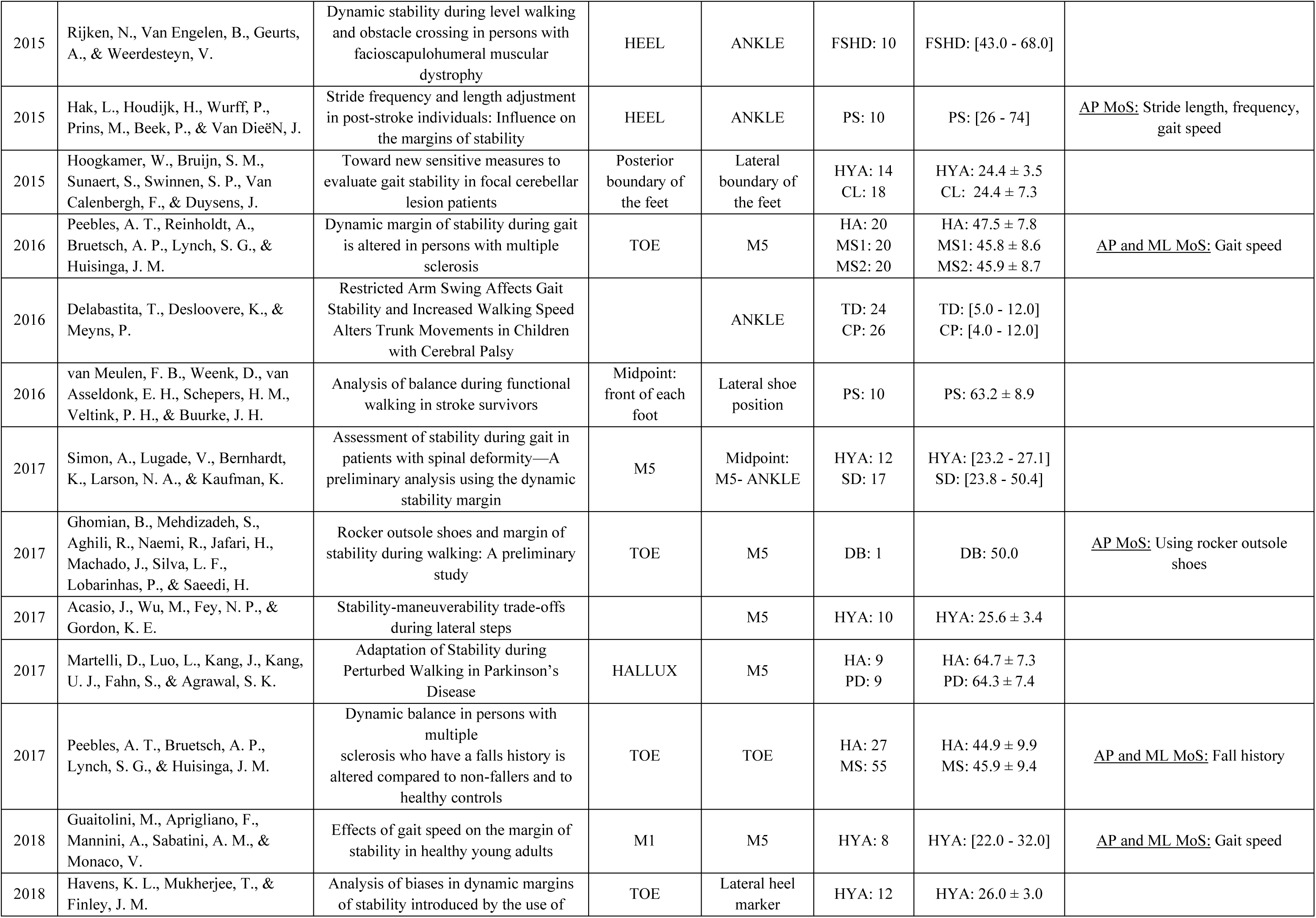
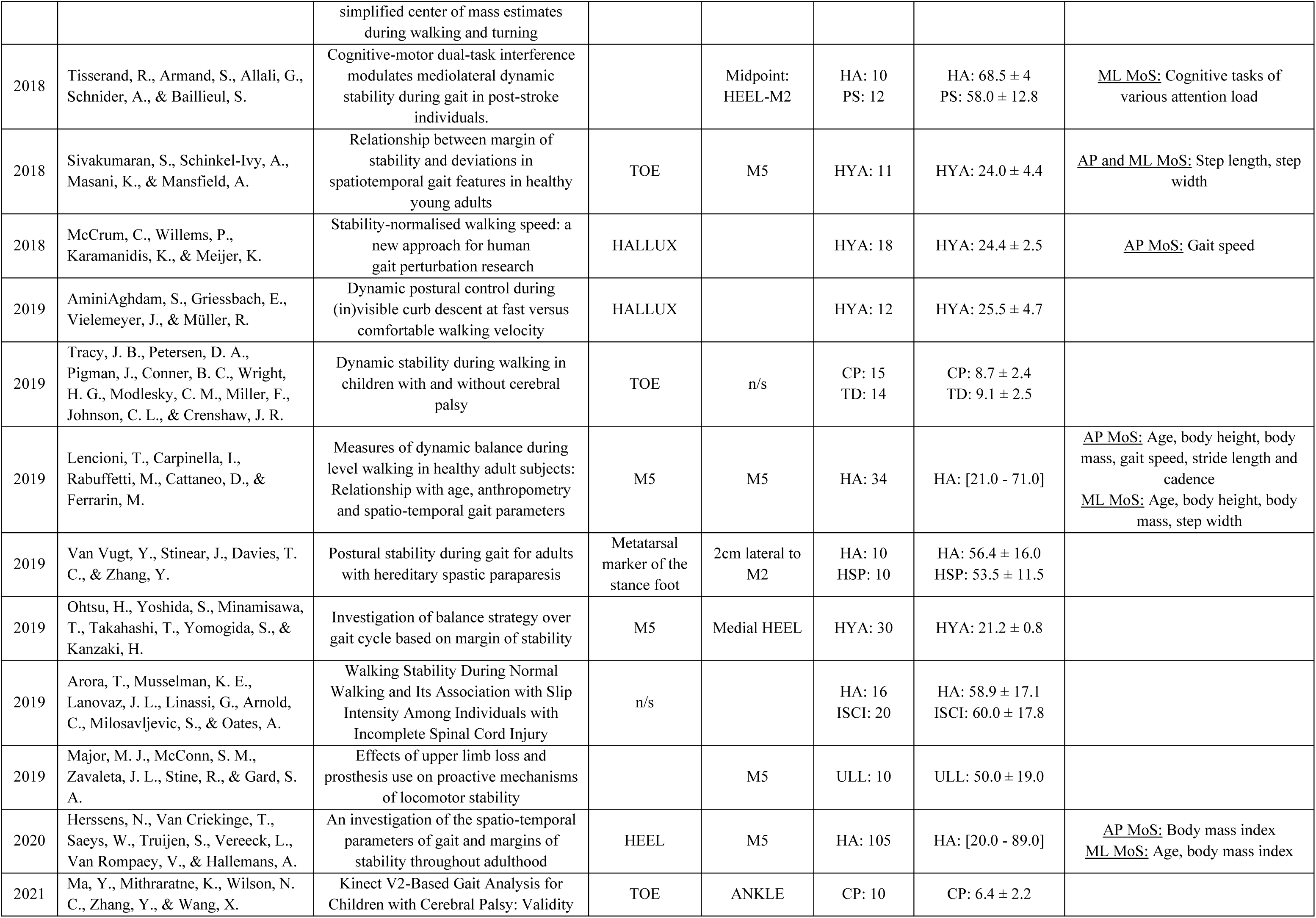
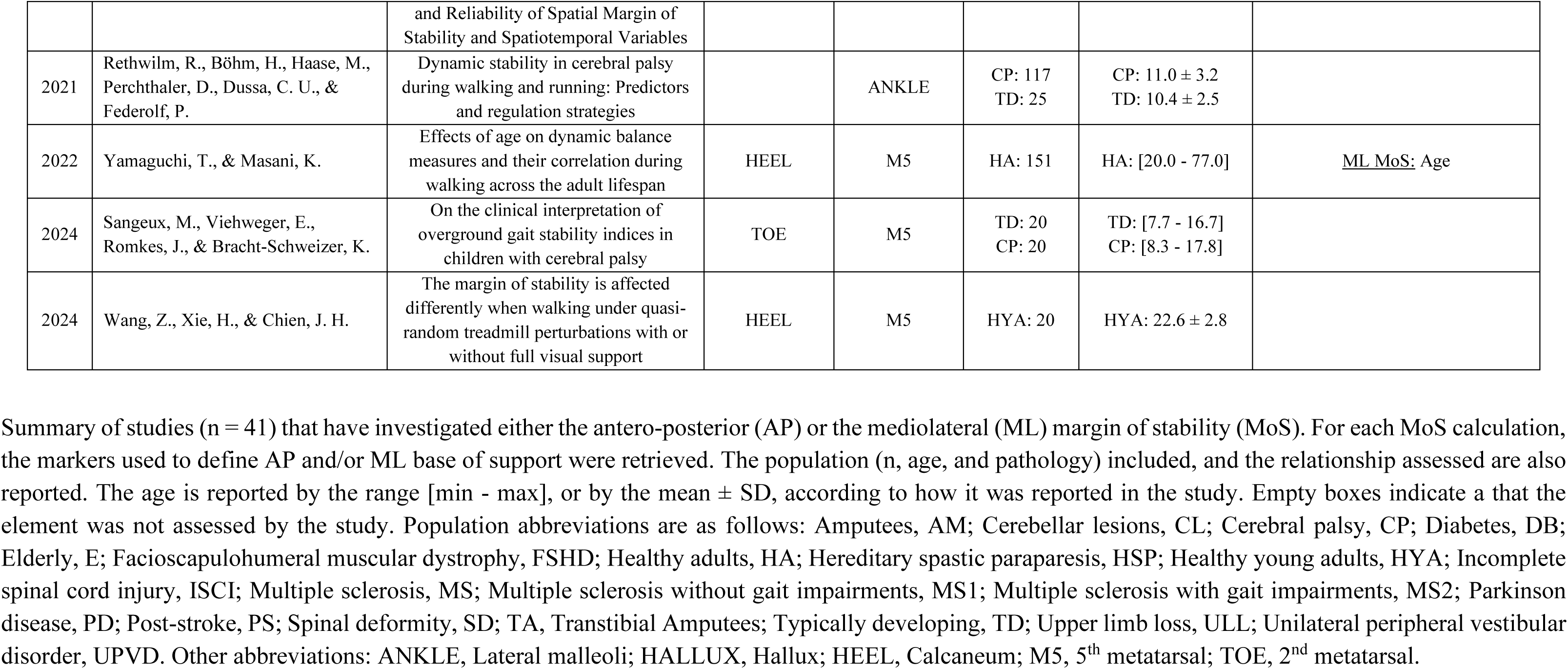
Literature review on the different approaches used to calculate the margin of stability.

